# Crystal structure of the membrane anchoring domain of mycobacterial Wag31: a dimer-of-dimer suggests how a Wag31 filament might self-assemble

**DOI:** 10.1101/869594

**Authors:** Komal Choukate, Barnali Chaudhuri

**Author notes:** corresponding author, Address: G. N. Ramachandran protein center, CSIR Institute of Microbial Technology, Sector 39A, Chandigarh 160036, India.

## Abstract

Wag31, or DivIVA, is an essential protein and a drug target in human pathogen *Mycobacterium tuberculosis* that self-assembles at the negatively curved membrane surface to form a higher-order structural scaffold, maintains rod-shaped cellular morphology, and localizes key cell-wall synthesizing proteins at the pole for exclusive polar growth. We determined the crystal structure of N-terminal membrane anchoring domain of mycobacterial Wag31 at 2.3 Å resolution using molecular replacement method. Crystal packing analysis revealed a previously unseen dimer-of-dimer assembly state of N-terminal Wag31 with C_2_ point group symmetry, which is formed by antiparallel stacking of two coiled coil dimers. Size-exclusion column chromatography-coupled small angle solution X-ray scattering data showed a tetrameric form as a major assembly state of N-terminal Wag31 in solution, further supporting the crystal structure. Plausible models of linear self-assembling, and branching, of Wag31 filaments consistent with available data are suggested.

## Introduction

Sensing of membrane curvatures play critical roles in diverse physiological processes such as maintenance of cellular morphology, polar or hyphal growth in bacteria and endocytosis in eukaryotes (Cannon et al., 2017). Prominent examples of positive curvature sensing proteins include septins and BAR (Bin/amphiphysin/Rvs) domain proteins in eukaryotes and SpoVM in sporulating bacteria (Cannon et al., 2017). In gram positive bacteria, DivIVA recognizes negative, concave membrane curvature, and self-assembles at the cytoplasmic side of the pole and curved region of the cell division septum to form a structural scaffold (Edwards and Errington, 1997; Letek et al., 2008; Lenarcic et al., 2009; Ramamurthi and Losick, 2009; Kaval and Halbedel, 2012; Bach et al., 2014). DivIVA, which is a filamentous, coiled-coil protein, aids in the localizations of various target proteins at the pole for cell-wall growth and a variety of biological functions (Rudner and Losick, 2010; Laloux and Jacobs-Wagner, 2014; Halbedel and Lewis, 2019; Hammond et al., 2019).

DivIVA, also known as Wag31 and antigen 84, is essential in *mycobacteria*, and is required for its exclusive polar growth, morphology maintenance and other functions (Nguyen et al., 2007; Kang et al., 2008; Mukherjee et al., 2009; Jani et al., 2010; Plocinski et al., 2012; Ginda et al., 2013; Plocinska et al., 2014; Melzer et al., 2018). Furthermore, Wag31 is targeted by bactericidal amino-pyrimidine sulfonamides, though the mechanism of antibacterial action is not known (Boshoff, 2017; Singh et al., 2017). Wag31 play critical roles in regulating peptidoglycan biosynthesis and localizing many cell-wall synthesizing enzymes at the pole to support polar growth (Kang et al., 2008; Jani et al., 2010; Meniche 2014; Xu et al., 2014). Wag31 is phosphorylated at a single site (T73), enabling it to be better at localization at the pole and at regulation of peptidoglycan biosynthesis (Kang et al., 2008; Jani et al., 2010). A small membrane bound protein CwsA may assist Wag31 localization at the mycobacterial pole (Plocinski et al., 2012). While depletion of Wag31 leads to “rod to spherical cell” transition (Nguyen et al., 2007; Kang et al., 2008; Meniche et al., 2014), Wag31 is shown to contribute to restoration of rod shape in spherical cells (Melzer et al., 2018).

Oligomerization appears to be a recurring theme in the sensing of meso-scale (∼100 nm to 1 micron) curvatures by nano-scale protein subunits in several cases (Antonny, 2011; Cannon et al., 2017). A “molecular bridging of the curvature” by oligomers of DivIVA was suggested as a mechanism of DivIVA binding to concave membrane (Lenarcic et al., 2009). However, how DivIVA oligomerize is sparsely understood. Structural data on DivIVA is so far limited to only one study (Oliva et al., 2010). Model of a 30 nm long tetrameric form of DivIVA from *B. subtilis* (bsDivIVA) was proposed based on the crystal structures of individual N-terminal and C-terminal coiled coil domains of DivIVA (Oliva et al., 2010). Conserved phenyl alanine and arginine residue pairs at both ends of this bent, elongated tetrameric structure occupies the proposed sites of polar membrane tethering (Oliva et al., 2010). In addition, super-resolution microscopy revealed double-ring structures formed by self-assembled bsDivIVA near the division septum (Eswaramoorthy et al., 2011). Electron micrographs showing oligomerizations of bsDivIVA mutants to form ∼ 22 nm long “doggy-bone” shaped structures, and longer strings and networks *in vitro* were reported (Stahlberg et al., 2004).

Recently, we showed that the full-length mycobacterial Wag31 (tbWag31, P9WMU1, Rv2145c) form several microns long primarily linear polymers *in vitro*, with occasional bending or branching (Choukate *et al.*, 2019). Here, we report a crystal structure of the N-terminal membrane binding domain of tbWag31 (N-Wag31) at 2.3 Å resolution. Crystal packing analysis reveals a tetrameric form of N-Wag31, which is composed of two dimeric coiled coil domains. Accompanying size-exclusion column chromatography-coupled small angle X-ray scattering (SEC-SAXS) data further supports a tetrameric form of N-Wag31 in solution. Based on the available data, we proposed models of Wag31 self-assembling for linear, and branched, filament formation.

## Results

### Crystal structure of the N-terminal domain of Wag31

Mycobacterial Wag31 is a 260 residule long filament-forming protein containing two domains: an N-terminal lipid or membrane binding domain and a C-terminal domain that participates in polar protein localization. Crystal structure of the N-terminal domain of tbWag31, or N-Wag31, shows a parallel coiled-coil dimer composed of two chains, A and B, containing 2-60 residues and additional residues from the poly-histidine tag regions (figure 1, see figure S1 and methods section). The two chains of N-Wag31 are similar to each other, with root mean square deviation or RMSD of 0.65 Å for 59 C*α* atoms (calculated using CLICK, Nguyen et al., 2011). The N-terminal segment of N-Wag31 contains a short helical turn (H1 helix) and loop followed by a sharp turn joining the coiled coil helix (H2 helix). The loop region is intertwined in the N-Wag31 dimer, as observed for the N-terminal domains of its structural homologs bsDivIVA (N-bsDivIVA), and GpsB (Oliva et al., 2010; Halbedel and Lewis, 2019; Cleverley et al., 2019). Circular dichroism data obtained from N-Wag31 supported an alpha-helix rich N-Wag31 structure in solution, with an estimated helical content of ∼ 60 % (figure 1; calculated using K2D; Andrade et al., 1993). The average coiled coil pitch is ∼ 171 Å for 26-60 residues of N-Wag31 (calculated using TWISTER, Strelkov and Burkhard, 2002).

**Figure 1.**
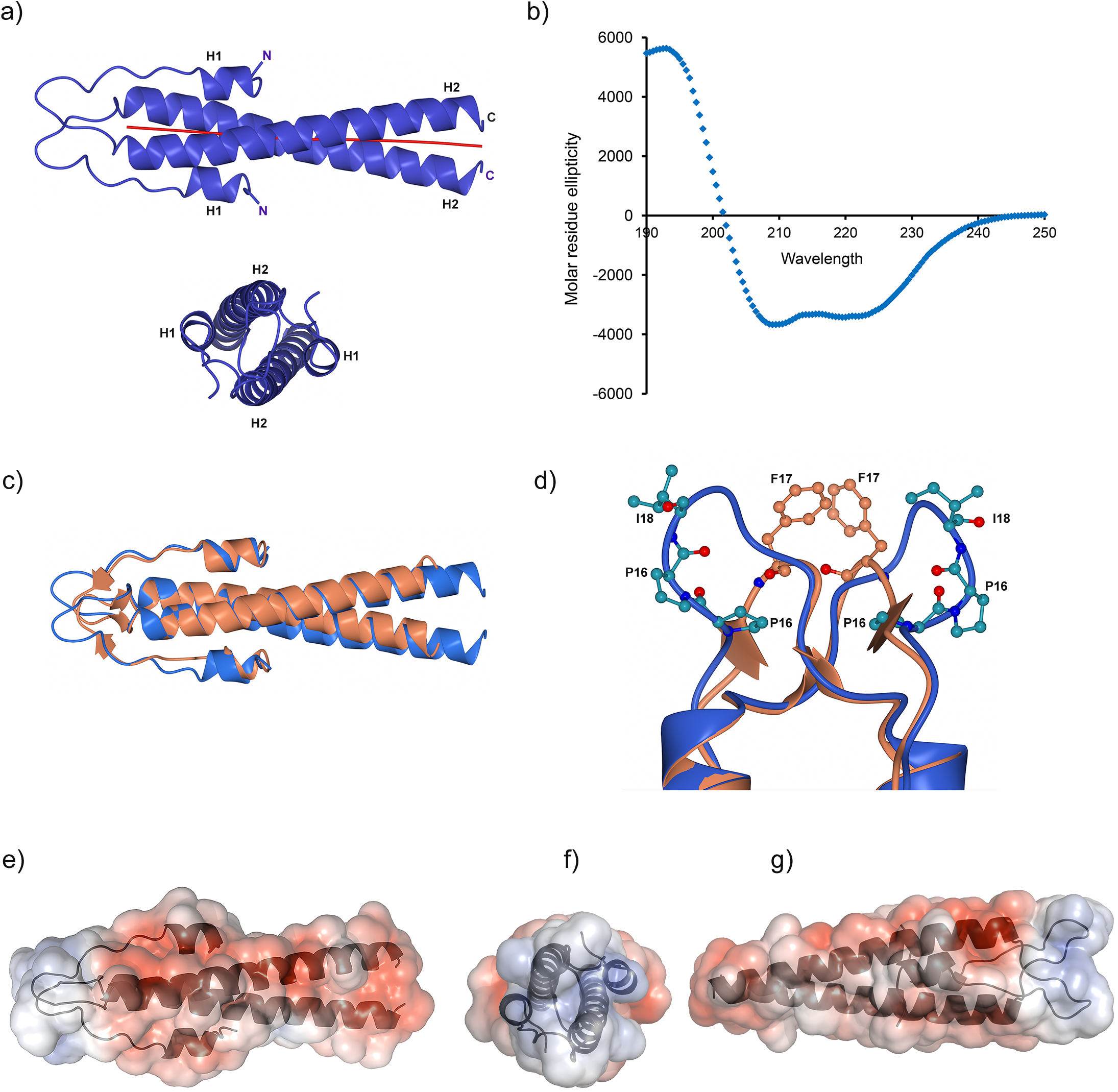
Crystal structure of N-Wag31. (a) Two nearly-orthogonal views of N-Wag31 (2-60 residues) dimer as blue ribbons are shown. The calculated local coiled coil axes are shown as red jointed lines. (b) CD profile (molar residue ellipticity *versus* wavelength) of N-Wag31. (c) Superimposed structures of N-Wag31 (blue) and N-bsDivIVA (coral) are shown. (d) The intertwined loop region in N-Wag31 (blue) containing the P_16_P_17_I_18_ sequence (ball and stick, with carbon atoms in light blue, oxygen in red and nitrogen in blue), and equivalent region in N-bsDivIVA (coral) containing the stacked F17 residues (ball and stick, with carbon atoms in coral, oxygen in red and nitrogen in blue), are shown. (e-g) Three views of the tetrameric N-Wag31 (grey) with electrostatic potential mapped into the water-accessible surface (water radius 1.4 Å) are shown. Electrostatic potential was computed using APBS with AMBER charges and suggested parameters (Baker et al., 2001). Figures were prepared using Pymol (The PyMOL Molecular Graphics System, Schrodinger, LLC.) and CCP4MG from the CCP4 suite (Winn et al., 2011).

N-wag31 structure is similar to the structure of its homolog N-bsDivIVA, with which it shares ∼ 42 % sequence identity, with 1.6 Å RMSD for 56 C*α* atoms in a chain (computed using Dali, Holm, 2019). However, the two structures differ significantly in the intertwined loop region that houses the lipid binding site (figure 1). In the N-bsDivIVA structure, two spatially adjacent phenyl alanine residues (phe17) from the intertwined loops of two subunits in the coiled coil dimer were shown to participate in lipid binding (figure 1; Oliva et al., 2010; Halbedel and Lewis, 2019). Arg18 side chains adjacent to these phe17 residues in N-bsDivIVA dimer likely interact with the negatively charged polar head groups of the membrane phospholipids (Oliva et al., 2010; Killian and von Heijine, 2000), which is conserved (arg21) in N-Wag31. The putative lipid-binding region in N-Wag31 houses a conserved “P_16_P_17_I_18_” motif instead of phe17 (figure 1). Exposed, non-polar ile18 at the tip of the intertwined loop is a candidate for direct interaction with membrane lipid. However, distance between the C*α* atoms of the two ile18 residues in N-Wag31 are ∼ 15 Å. In comparison, the C*α* atoms of lipid binding phe17 residues in N-bsDivIVA are separated by ∼ 5 Å, with their side-chain aromatic rings stacked (figure 1). A conformational change may possibly bring the ile18 hydrophobic side-chains together in N-Wag31 to form a lipid interacting patch. On the other hand, presence of two proline residues before ile18 makes this region structurally rigid. Thus, the putative lipid binding region is quite different in N-Wag31 than in N-bsDivIVA.

Mapping the Poisson-Boltzmann electrostatic potential into the solvent accessible surface revealed that the surface of the lipid binding N-Wag31 dimer is highly polar (figure 1). Note that, surface residues with poor electron density in N-Wag31 were modelled as suitable rotamers from the penultimate library (Lovell et al., 2000). N-Wag31 sequence contains 14 negatively charged and six positively charged residues. Surface of the homologous N-bsDivIVA, with several charged residues, is similarly polar (Oliva et al., 2010). This is not surprising, as charged amino acids are more common in coiled coils than other proteins (Surkont and Pereira-Lal, 2015). The electrostatic potential calculation shows a positively charged patch at the intertwined loop region of N-Wag31, that likely defines the conserved membrane-associating region in Wag31/DivIVA family of proteins.

### Crystal packing analysis suggests a ‘dimer-of-dimer’ organization of N-Wag31

Coiled coils can assemble in a number of ways to form higher order assemblies (Moutevelis and Woolfson, 2009). In order to learn about assembling of N-Wag31, we performed crystal packing analysis. It appears that the dimeric coiled coil domain of N-Wag31 combines with an adjacent crystallographic two-fold symmetry related dimer to form a tetramer with C_2_ point group symmetry (figure 2). Isologous interface in N-Wag31 was assigned using PDBePISA server (Krissinel and Henrick, 2007). The tetramer buries ∼ 396.0 Å^2^ of water accessible surface area upon assembling, with the coiled coil-forming H2 helices of the two B-chains with complimentary surface shapes stacking against each other (figure 2). The two stacked H2 helices are in near-antiparallel orientation, with ∼ 176° angle between the two local helix axes (26-41 residues). In addition, a PEG molecule was found buried at one side of this assembly interface, covering a rather small, ∼ 77 Å^2^, part of the accessible surface area of the N-terminal H1 helix (figure 2). Note that formation of a tetrameric form of N-Wag31 in the absence of PEG is supported by solution SAXS data, which is described in the next section.

**Figure 2.**
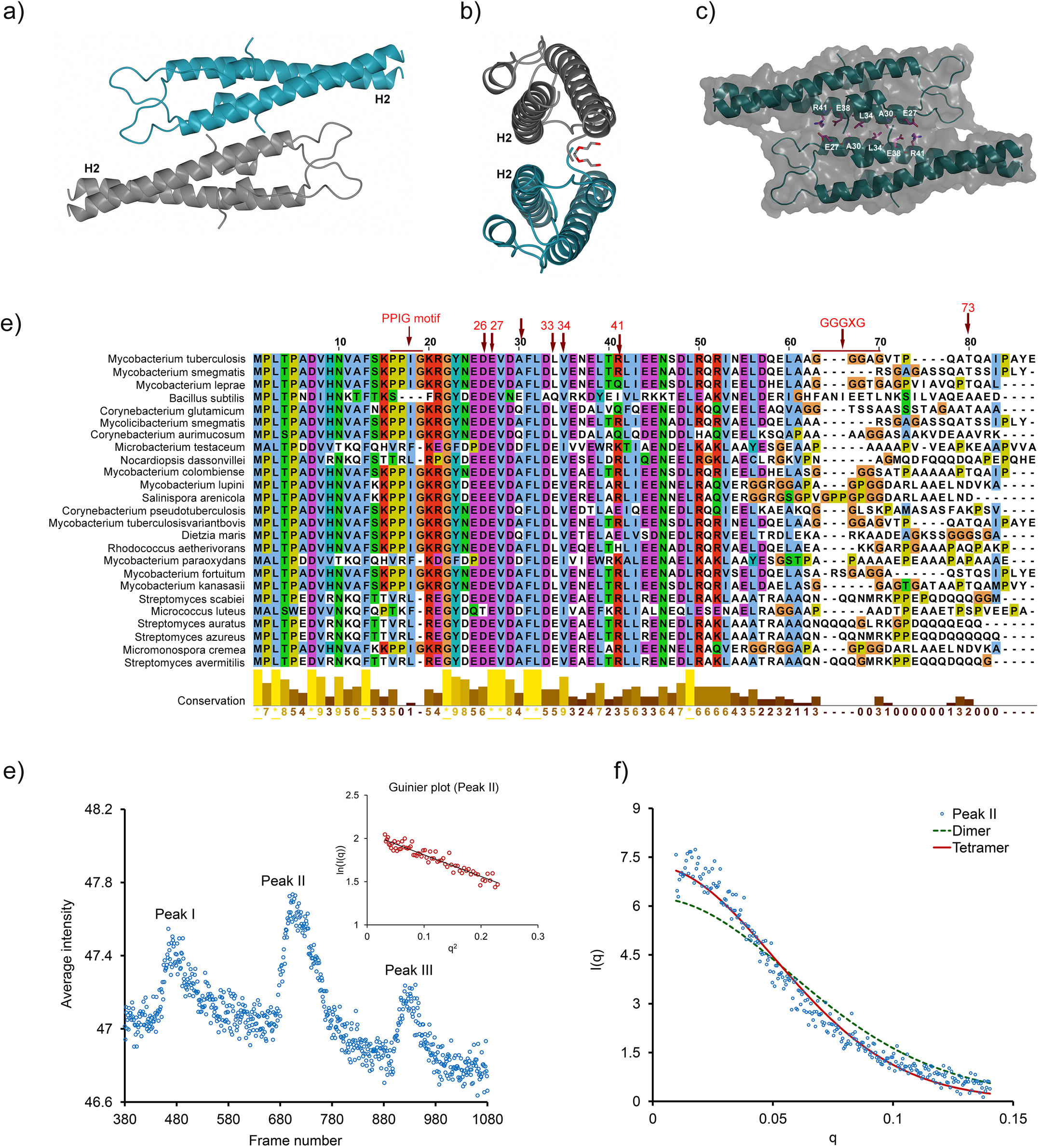
Oligomeric organization of N-Wag31. (a-b) Two views of the dimer-of dimer of N-Wag31 related by crystallographic 2-fold symmetry are shown as ribbons (cyan and grey). The interface-forming H2 helices are marked. A PGE molecule bound to N-Wag31 is shown as sticks in (b). (c) Tetrameric N-Wag31 is shown as ribbon (cyan) within the molecular surface (grey) of the tetramer. Side-chains of the interface forming residues R41, E27, L34, A30 and E38 are shown in ball-and-stick (carbon in magenta, oxygen in red and nitrogen in blue). (d) Multiple sequence alignment of the N-term region of tbWag31 (numbering for tbWag31 in red) and homologs, with the degree of conservation, is shown (prepared in Jalview, Waterhouse et al., 2009). (e) SAXS elution profile (average intensity *versus* frame number obtained from CHROMIXS, Franke et al., 2017) of N-Wag31. Guinier plot (lnI(q) *versus* q^2^, I is intensity and q is momentum transfer in nm^−1^) for peak II is shown in the inset. (f) Scattering profile of peak II data (I *versus* q in Å^−1^). Computed scattering profiles of dimeric and tetrameric N-Wag31 (calculated using CRYSOL with suggested parameters, Franke et al., 2017) are shown fitted to the experimental data.

Size of the buried surface area is a critical quantity that helps in distinguishing between crystal contacts and biological interfaces (∼370 – 4750 Å^2^ for a homodimer, Henrick and Thornton, 1998; Krissinel and Henrick, 2007). Buried surface area at the crystallographic 2-fold related interface in N-Wag3 is small (∼ 396.0 Å^2^). This crystallographic interface is lined by residues that are conserved in many actinobacteria, such as asp26, glu27, ala30, leu34 and arg41 (figure 2). Center of this interface is formed by buried non-polar leu34 and ala30. Two salt-bridges are formed between the side-chains of glu27 and arg41 residues at this interface on both sides of the central non-polar region (figure 2).

### Solution scattering supports a ‘dimer-of-dimer’ organization of N-Wag31

In order to determine the solution assembly states of N-Wag31, we performed SEC-SAXS experiment. Size-exclusion column elution profile of N-Wag31 revealed presence of multiple assembly states eluted as separate peaks, numbered from left to right as peaks I, II and III, respectively (figure 2). The early-eluting peak I suggests presence of a large aggregate. The averaged SAXS data obtained from peak III was too weak for analysis. SAXS data analysis suggests that the peak II region contains a tetrameric form of N-Wag31, which has a molecular mass of about ∼ 8.6 kDa/monomer, with a predicted mass of about 34.6 kDa based on Bayesian inference method (Hajizadeh et al., 2018). The −9 to 0 residues of the chain A of N-Wag31 containing a near-complete poly-histidine tag was used to build equivalent region of the chain B of the dimer, and the tetramer, using non-crystallographic symmetry operations, for all SAXS calculations. However, this tag region might be floppy in solution. Radius of gyration (R_g_) obtained from the Guinier analysis of averaged SAXS data from peak II was 27.2 Å (q.Rg ≤ 1.3, figure 2), while calculated R_g_ from dimeric/tetrameric N-Wag31 co-ordinates were about 20.9 and 24.2 Å, respectively. Chi-square values for fits between the theoretical scattering intensities derived from the tetrameric and the dimeric N-Wag31 models with experimental averaged scattering intensity from peak II were 1.45 and 4.0, respectively, further supporting the tetrameric structure (figure 2). As the averaged SAXS data was quite noisy in the high angle region and inadequate for further analysis, shape reconstructions were not performed. To summarize, solution SEC-SAXS data supports a tetrameric assembly of N-Wag31, which is consistent with the crystal structure.

### A suggested model of Wag31 filament formation

A protein subunit can be arranged to from a filament in a few ways. It could polymerize by pure translational repeat along a certain direction, or it can polymerize by a combination of rotation and translation along a helical axis. Alternatively, a protein subunit can form a filament by utilizing two-fold symmetry operations. For a protein with an N-terminal domain N and a C-terminal domain C, two such theoretical arrangements will be -NC-NC-NC-NC- or -NC-CN-NC-CN-NC- (figure 3). In the latter case, N-N and C-C inter-subunit interfaces related by two-fold symmetry will be formed (figure 3). Wag31 appears to form a linear filament using the second option involving two-fold symmetry. The N-Wag31 forms a tetramer with the C-terminal regions extending in opposite directions, which is compatible with linear filament formation (figures 2 and 3). Structure of the C-terminal domain of tbWag31 is currently not available. The C-terminal domain of bsDivIVA was shown to form a dimer of dimer, with a central four-helix bundle region (figure 3, Oliva et al., 2010). A combination of the two such N-terminal and C-terminal dimer-of-dimers can be used to build a linear filament (figure 3). Such linear filaments, with sporadic branching, were reported for full-length tbWag31 earlier (Choukate et al., 2019). Furthermore, the two-fold symmetry related interfaces in the N-Wag31 suggest a natural way for lateral or side-way association of dimeric N-Wag31 units to form a hexamer and other higher order oligomers (figure 3). Such a theoretical hexameric association of N-Wag31 can account for branching or bifurcations in a Wag31 filament, and can be exploited for protein-based scaffold design (figure 3).

**Figure 3:**
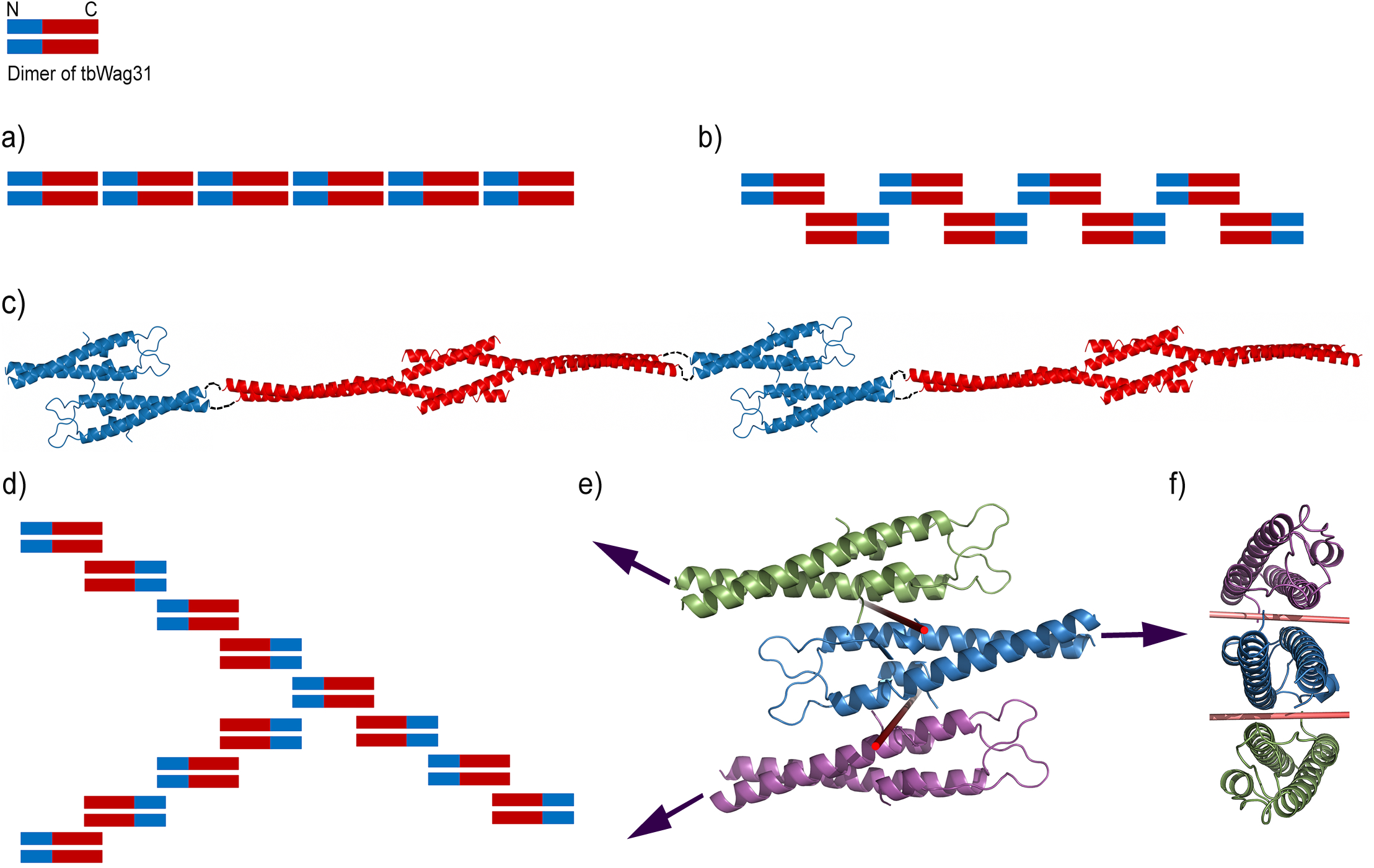
A proposed model of Wag31 filament assembling. (a-b) Two theoretically alternative possible modes of self-assembling of Wag31, using (a) pure translational repeat, and (b) two-fold symmetry related repeat, are shown. (c) Ribbon diagrams of the tetramers of N-terminal N-Wag31 (blue) and C-terminal domain of bsDivIVA (red) are shown, joined by linker region as dashed lines, as a theoretical filament. The linker region is ∼20 residue long in bsDivIVA, and ∼100 residue long in tbWag31. (d) A theoretically possible mode of 3-way branching in tbWag31 filament. (e-f) Two views of a theoretical hexamer composed of three dimers of N-Wag31 (ribbons in green, blue and magenta). The two-fold axes of rotation are shown as red rods.

## Discussion

We report crystal structure of the lipid binding domain of mycobacterial Wag31. The positively charged, putative lipid binding region in this structure is strikingly different from the lipid binding region of its structural homolog N-bsDivIVA, suggesting that tbWag31 may use a different mode of lipid binding than bsDivIVA (figure 1). This site in N-Wag31 may form a binding site for CwsA that aids in polar localization of Wag31 (Plocinski et al., 2012). Crystal structure of N-Wag31 reveals a tetrameric form of N-Wag31, which is further supported by solution SAXS data. Considering that tbWag31 is a high confidence drug target (Singh et al., 2017; Boshoff, 2017), our structural data can be exploited for designing inhibitors and for designing mutations to probe tbWag31 function *in vivo*.

Crystal structure of N-Wag31 that comprises of the first 60 residues of tbWag31 raises question about the possible structural role of phosphorylated T73 residue in full-length tbWag31. The T73 residue is in a putative intrinsically unstructured region adjacent to the N-terminal domain (Choukate et al., 2019), and proximal to several glycine residues, such as G_63_G_64_G_65_X_66_G_67_ (figure 2). Small and flexibility-imparting glycine residues are typically rare in coiled coils (Surkont and Pereira-Lal, 2015). Within the classical heptad repeat “abcdefg” in a coiled coil, ‘a’ and ‘d’ regions are buried inside were ‘g’ and ‘e’ regions make ionic interactions between the helices to stabilize the coiled coil assembly. If the coiled coil heptad assignment (made by TWISTER; Strelkov and Burkhard, 2002) is theoretically extended beyond the last residue (leu60, assigned ‘a’ position) observed in the crystal structure, the 73^rd^ residue will be in ‘g’ position assuming coiled-coil continuity, and could be critical for providing a stabilizing salt-link for coiled coil continuation in the phosphorylated form of tbWag31 in an otherwise flexible region.

The tetrameric form of N-Wag31 naturally explained linear, and branched, assembly formation of tbWag31 (figure 3). Full-length tbWag31 forms linear filament, which is at times branched (Choukate et al., 2019). Filaments, networks and doublering structures formed by bsDivIVa were reported earlier (Stahlberg et al., 2004; Eswaramoorthy et al., 2011). Our structural data in combination with previously published data suggest plausible models for such filament formation. These results, in addition to aiding in understanding Wag31 function, and aiding antibiotic discovery efforts, could be exploited for protein assembly design purpose.

### Experimental method

#### Recombinant Protein Expression

The N-terminal domain of Wag31 (2-60 aa with an N-terminal tag containing poly-histidine residues, synthesized by Genescript) construct in pET15b vector was expressed in *Escherichia coli* BL21 (DE3) cells. A primary culture of 20 ml volume was started at 37° C in LB broth containing 100 mg/ml ampicillin and incubated overnight. This primary culture was transferred and grown in 2 liters of LB broth at 37° C until the OD_600_ of 0.6 was reached. Next, recombinant protein expression was induced by adding 1 mM IPTG and cells were incubated at 25° C for 16 hrs. The cells were harvested and stored at −80° C for further use.

#### Protein Purification

The frozen cell pellets were resuspended in lysis buffer containing 20 mM Tris HCl pH 7.0, 150 mM NaCl with 0.2 % Triton-X 100 and lysed by sonication. The cell lysate was loaded onto Nickel IMAC sepharose fast flow affinity column and bound protein was eluted with a buffer containing 50 mM Tris.HCl at pH 7.0, 150 mM NaCl and 500 mM imidazole. The fractions of interest were pooled, concentrated and further purified by size-exclusion chromatography using a HiLoad 16/60 Superdex 200pg column in following buffer: 50 mM Tris.HCl at pH 7.0 and 150 mM NaCl. Freshly purified N-Wag31 protein was concentrated to 10 mg/ml for crystallization experiments.

#### CD spectro-polarimetry

For secondary structure estimation, CD spectra of purified N-Wag31 at ∼ 1 mg/ml concentration was measured at 25° C using an in-house Jasco J815 spectropolarimeter. Three CD spectra scans were averaged and plotted as molar residue ellipticity *versus* wavelength.

#### Crystallization, Data collection, Structure Determination and Refinement

Crystals of N-Wag31 were grown by both hanging and sitting drop vapour diffusion methods at 19° C in 0.2 M Magnesium chloride, 0.1 M sodium cacodylate at pH 6.5 and 50% (v/v) polyethylene glycol 200. These crystals were very small and recalcitrant to growth. Crystals were flash-frozen in liquid nitrogen without any additional cryo-protectant and transported to ESRF, France. X-ray diffraction data from an N-Wag31 crystal was collected at the microfocus beamline ID30A-3 (MASSIF-3) at ESRF, which is suitable for such small crystals. Only one useful dataset was obtained after trying 37 crystals. Data collection was done at 100 K, using an Eiger X 4M detector (DECTRIS Ltd.). Data indexing, processing, merging and scaling were performed using XDS (Kabsch, 2010) and the CCP4 suite of software (Winn et al., 2011).

The crystal structure of N-Wag31 was determined using molecular replacement method (MOLREP, Vagin and Teplyakov, 1997). The sole available crystal structure of its homolog bsDivIVA (PDB code 2wuj; Oliva et al., 2010) was used as a search model in a dimeric form, after converting it to a poly-alanine model. Following rigid body minimization of the best solution, positional refinements were performed using Refmac (Murshudov et al., 1997) with NCS restraints. R-free set was used to monitor the progress of refinement. Individual isotropic B-factors were refined. Model rebuilding was performed using Coot (Emsley et al., 2010). SigmaA weighted difference maps and composite omit maps were used for model building. In the latter stages of refinement, simulated annealing were performed in Phenix (Adams et al., 2010). NCS restraints were released during the final rounds of the refinement. In addition to two protein chains and water molecules, the asymmetric unit contains a triethylene glycol (PGE). Molprobity was used for structure validation (Chen et al., 2010).

N-Wag31 contains two chains, A and B, in the asymmetric unit. In addition to residues 1-60, chain A contains a large, contiguous segment of the poly-histidine tag region (−9 to 0 residues). Chain B contains 2-60 residues. An isolated tripeptide interacting with a neighboring molecule, tentatively containing “HHE” sequence, was assigned to chain B. However, as this tripeptide segment is far from the rest of the B chain, an out-of-register may not be ruled out. Electron density at and around the loop region containing 17-20 residues, especially in chain A, was rather poor. Main-chain conformation for this loop region was traced and built in the electron density map for chain B and rebuilt in the same way for chain A. Side-chain densities were poor for several residues around this loop region and at the C-terminal end of N-wag31, which is reflected in higher than average temperature factors for these side-chains. All these side-chains were modelled using suitable rotamers from the coot rotamer library (Lovell et al., 2000). Loss of side-chain density could be due to radiation damage caused by X-ray at the microfocus beam-line. Data collection and refinement statistics are summarized in table 1. Figures were prepared using Pymol (The PyMOL Molecular Graphics System, Schrodinger, LLC.) and CCP4MG from the CCP4 suite (Winn et al., 2011).

**Table 1.** Crystal diffraction and model refinement statistics (generated using the table 1 option of Phenix, Adams et al., 2010). Statistics for the highest-resolution shell are shown in parentheses.

| **(a) Diffraction data and refinement statistics** | **N-Wag31** |
| --- | --- |
| Experimental station | ID30A-3, ESRF |
| Wavelength (Å) | 0.97 |
| Resolution range (Å) | 30.1 - 2.3 (2.4 - 2.3) |
| Space group | C 1 2 1 |
| Unit cell (a, b, c in Å and $\alpha$, $\beta$, $\gamma$ °) | 44.9, 53.8, 61.2, 90, 100.8, 90 |
| Total reflections | 77318 (5929) |
| Unique reflections | 6374 (592) |
| Multiplicity | 12.1 (9.9) |
| Completeness (%) | 98.6 (92.2) |
| Mean I/sigma(I) | 7.45 (2.2) |
| Wilson B-factor (Å<sup>2</sup>) | 21.75 |
| R-merge | 0.235 (1.054) |
| R-meas | 0.245 (1.112) |
| R-pim | 0.070 (0.345) |
| CC1/2 | 0.99 (0.76) |
| CC* | 0.99 (0.93) |
| Reflections used in refinement | 6363 (592) |
| Reflections used for R-free | 353 (38) |
| R-work | 0.175 (0.180) |
| R-free | 0.225 (0.245) |
| Number of non-hydrogen atoms | 1137 |
| - Macromolecules | 1091 |
| - PGE | 10 |
| - Water | 36 |
| Protein residues | 132 |
| RMS (bonds, Å) | 0.010 |
| RMS (angles, °) | 1.05 |
| Ramachandran favored (%) | 98.4 |
| Ramachandran allowed (%) | 1.6 |
| Ramachandran outliers (%) | 0.00 |
| Rotamer outliers (%) | 0.8 |
| Average B-factor (Å<sup>2</sup>) | 32.6 |
| - Macromolecules | 32.6 |
| - PGE | 35.5 |
| - Water | 31.3 |

### Size-Exclusion chromatography coupled with SAXS experiment

SEC-SAXS experiment was performed at the BM29 beam-line at ESRF, using a Shimadzu HPLC system. The purified N-Wag31 (4.4 mg/ml) in a buffer containing 20 mM Tris pH 7.5, 150 mM NaCl and 10% glycerol was used for SAXS experiments. The protein samples were frozen under liquid nitrogen and transported to synchrotron site, where it was thawed on ice prior to the experiment. 30 micro liter of protein sample was injected to an Agilent Bio-SEC-3 column. SEC-SAXS data was collected at ∼ 1 Å X-ray wavelength, with 1 s/frame exposure and 2.8 m detector distance, using a Pilatus detector. SEC-SAXS data analysis was performed using the ATSAS suite of software (Franke et al., 2017).

## Data availability

Protein structure coordinates and structures factors are deposited to protein data bank (PDB, www.rcsb.org), with accession code 6LFA. SAXS data for peak II is available from SASBDB (sasbdb.org), with accession code xxxx (deposition draft ID 2063).

## Acknowledgements

Authors would like to thank CSIR for intramural research support and fellowship to KC. We thank Department of Biotechnology, and ESRF-DBT access program administered by Regional Center of Biotechnology, for supporting X-ray diffraction and SEC-SAXS data collection at ESRF. We thank Dr. Nishant Varshney and staffs at BM29 and ID30 beamlines at ESRF for assistance during data collection. We thank Prof. Jan Löwe from MRC, Cambridge, for providing the coordinates of the C-terminal domain of bsDivIVA.

---

**Figure S1.**
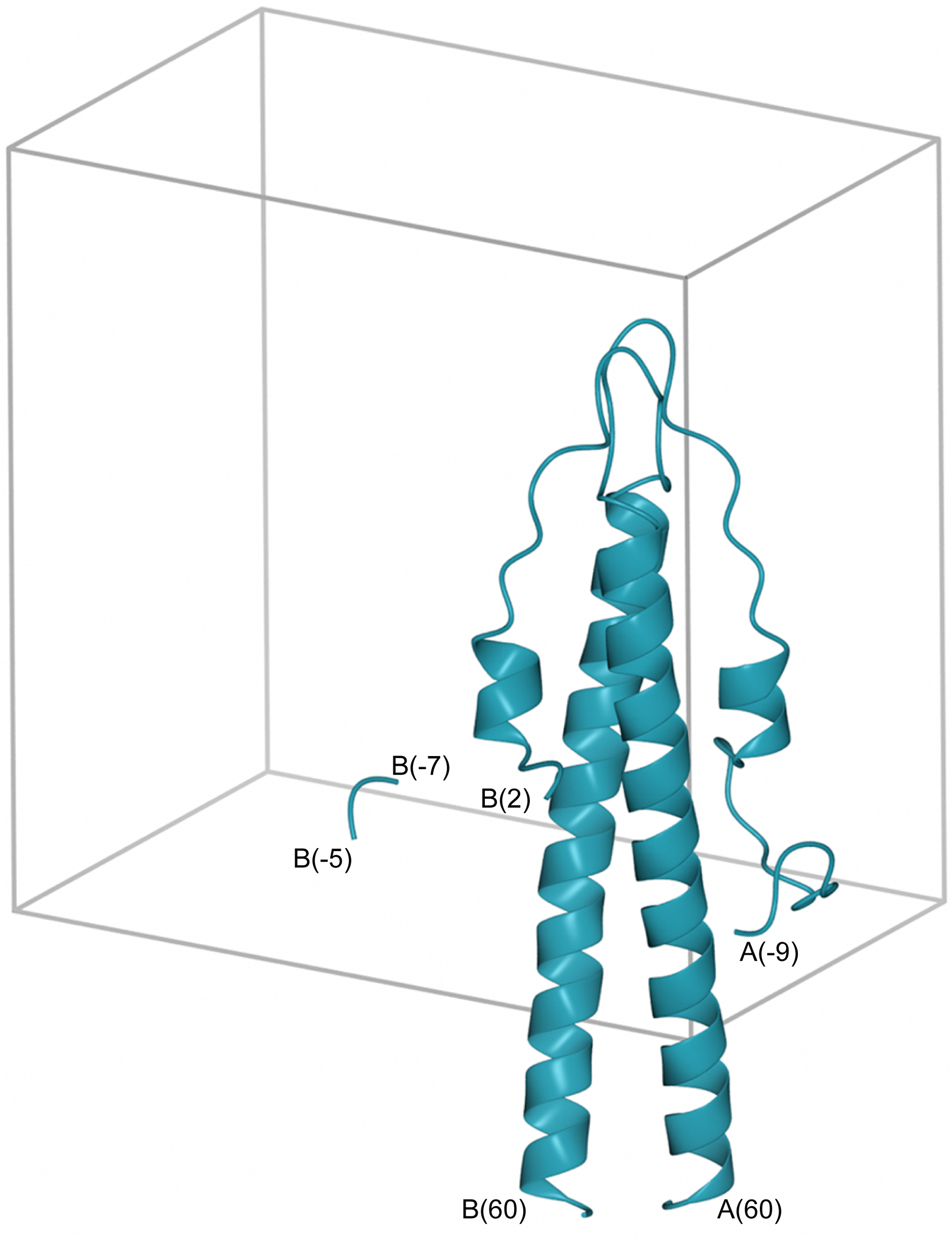
Structure of N-Wag31. The monoclinic unit cell is shown as a box. N-Wag31 dimer within the asymmetric unit, including the poly-histidine tag regions (see methods section), is shown as a cartoon.

